# Human Airway Mucociliary Tissue Cultures Chronically Exposed to E-Cigarette Vapors Exhibit Altered Cell Populations and Increased Secretion of Immunomodulatory Cytokines

**DOI:** 10.1101/2022.12.12.520128

**Authors:** Vincent J. Manna, Shannon Dwyer, Vanessa Pizutelli, Salvatore J. Caradonna

**Author notes:** Contact, Department of Molecular Biology, Rowan University School of Osteopathic Medicine, 42 East Laurel Road, Suite 2200, P.O. Box 1011, Stratford, New Jersey 08084.

## Abstract

Vape-pens or electronic cigarettes (e-cigarettes) are handheld battery powered devices that use a vape-liquid to produce a vapor that is inhaled by the user. The active ingredients in commercial vape-liquids are commonly nicotine, tetrahydrocannabinol or cannabidiol. A consequence of the rise in e-cigarette usage was the 2019 emergence of a vaping-induced respiratory disease denoted ‘e-cigarette or vaping use-associated lung injury’ (EVALI). One of the suspected causes of EVALI is Vitamin E Acetate (VEA), which was found to be a diluent in certain illicit tetrahydrocannabinol vape-pens, whereas nicotine is commonly diluted in equal parts propylene glycol and vegetable glycerin (PG:VG). The prevalent use of e-cigarettes by both adult and young adult populations and the emergence of a novel illness has made understanding how e-cigarette vapors affect our respiratory tissues a public health concern. We have designed and produced a simple device that can operate commercial e-cigarettes and deliver the vapor to a chamber containing a standard cell culture multi-well plate. Here we utilize our device to investigate how human airway mucociliary tissue cultures respond after chronic exposure to vapors produced from either PG:VG or VEA. We note several differences between how PG:VG and VEA vapors interact with and alter airway tissue cultures and suggest potential mechanisms for how VEA-vapors can exacerbate EVALI symptoms. Our device combined with primary human airway tissue cultures make an economical and compact model system that allows for animal-free investigations into the acute and chronic consequences of e-cigarette vapors on primary respiratory cells.

## INTRODUCTION

Electronic cigarettes (e-cigarettes) and vape-pens are handheld battery-powered devices that contain a vape-liquid which is used to produce a vapor that is inhaled by the user. Vapor is produced by passing a current through a wire wrapped around a wick soaked in vape-liquid which then combusts to form the vapor that is inhaled [1]. The major active ingredients in commercial vape-liquids vary from nicotine to tetrahydrocannabinol (THC) or cannabidiol (CBD). E-cigarettes attract some traditional smokers as a safer alternative to smoking, but many users are teens and young adults. A CDC study concluded that from 2011 to 2018 e-cigarette use in high school students increased from 1.5% to 20.8% [2]. The global vape industry has a market size that was valued at 12.41 billion in 2019 and projected to rise over 20% within the next 5 years [3]. These statistics demonstrate that the sale and usage of e-cigarettes and other vape-products is growing rapidly.

One unforeseen byproduct of the booming vape industry was the June 2019 emergence of an illness that is linked to the use e-cigarettes and other vape-pen products. The Centers for Disease Control and Prevention (CDC) has termed the non-viral, non-bacterial, pneumonia-like illness “e-cigarette or vaping use-associated lung injury” (EVALI). The symptoms of EVALI include shortness of breath, fever and chills, cough, vomiting, diarrhea, headache, dizziness, rapid heart rate and chest pain. Since the initial emergence, EVALI has resulted in 2,800 hospitalized patients and 68 deaths in the US [4] [5]. Investigations led by the CDC into the cause of EVALI have highlighted vitamin E acetate (VEA) as a prime suspect [6]. VEA, an FDA approved substance that has been deemed safe for consumption and topical application, is added to vaping liquid as an additional solvent to increase total volume. It serves as what is commonly known as a “cutting agent”. A study performed by the CDC compared the presence of VEA in bronchoalveolar lavage (BAL) fluid in patients with EVALI to healthy participants. Of the 51 EVALI patient samples, 94% contained VEA, while none was detected in the control samples. The CDC has since concluded that VEA is not a suitable solvent for e-cigarettes and suggests that manufacturers remove it from vape-liquids until more studied are performed to determine the safety of vaping VEA.[5] [6]. These findings highlighted the need for evaluating the safety of other vape-liquid additives in addition to VEA.

The most common base solvents used in vape-liquids are propylene glycol (PG) and vegetable glycerin (VG) because of their FDA approval as safe for consumption and topical application in cosmetics as well as their lipophilic solvent properties. Although PG and VG are safe for consumption and topical applications, the vaping process is different and should be treated as such. Multiple chemical studies have revealed toxic by-products when vaping PG and VG liquids, including carbon monoxide, acetylene, methane, and aldehydes [7] [8] [9] [10]. In 2019 Song et al. performed a 4-week study where participants took 40 puffs per day from vape-pens containing only a 1:1 PG-VG mixture. The researchers concluded that the BAL fluid from the vape-user group had an increase in inflammatory cytokines and macrophages over control groups [11]. These studies further highlight the need for research into the safety of base solvents used in vape-liquids in addition to ‘cutting agents’ and active ingredients. A more systematic analysis of the consequences to the airway when vaping PG, VG and VEA will aide in navigating the future safety for vape users as well as bystanders worldwide.

One complication faced when researching the tissue consequences of e-cigarette vapor exposure is choosing an applicable model system. Although animal models have provided researchers with valuable information for generations, for ethical considerations it is always desirable to avoid animal models if an *in-vitro* model exists. Likewise, it is preferable to choose a human model over an animal model to minimize untranslatable findings due to molecular inconsistencies between humans and laboratory animals. The air-liquid interface (ALI) model system is widely used for generating human airway mucociliary tissue from primary airway cell cultures and fully differentiated ALI-tissue can be maintained in culture for multiple months which allows for long-term experiments. At the conclusion of experiments ALI tissue can be harvested for analyses including histology, immunohistochemistry, gene expression studies, western blotting, and other molecular techniques which provide a broad selection of data points for prospective studies.

Another difficulty in e-cigarette vapor research is the method of applying experimental vapors to the model system. Devices are available for use in delivery of volatile organic compounds and related environmental pollutants but are not designed for use with e-cigarettes [12] [13] [14]. Although a few companies offer experimental smoking machines designed for research into the effects of smoking cigarettes, these machines are large and expensive pieces of laboratory equipment that require considerable training for their operation and maintenance [13]. Our laboratory has taken advantage of 3D-Printing and microcontroller technology to develop a compact automated e-cigarette research device capable of operating e-cigarettes and delivering the vapor to a multi-well cell culture plate. Our laboratory combined the ALI model system with our vaping device to demonstrate the applicability for investigations in the study of inhaled/aerosolized toxins. By using the ALI tissue model system, we can focus on identifying cellular responses and consequences specific to airway mucociliary tissue, removing crosstalk or interference from immune cell or hormonal responses which may occur in animal-based model systems. In this study we compared the effects on differentiated ALI tissues chronically exposed to e-cigarette vapors generated from either PG:VG or VEA.

## METHODS

### Construction of Vaping Research Device

The vaping research device is 3D printed out of polylactic acid plastic and the components are controlled by an Arduino microcontroller. The device has a chamber that holds a standard 12-well cell culture plate, and a clamping mechanism that holds the experimental vape-pen (Figure 1). The vaping process is automated for maximum reproducibility and standardization between experimental vape-liquids. The process begins with a 15 second ‘inhale’ cycle, which utilizes a servo-driven button pushing mechanism and a fan-driven air pump to draw vapor out of the vape-pen and deliver it via holes centered above the transwell inserts for maximum vapor-to-tissue contact. After a 5 second pause the ‘exhale’ cycle begins when a second air-pump activates for 10 seconds that will clear the chamber of vapor and end the vaping process.

**Figure 1A:**
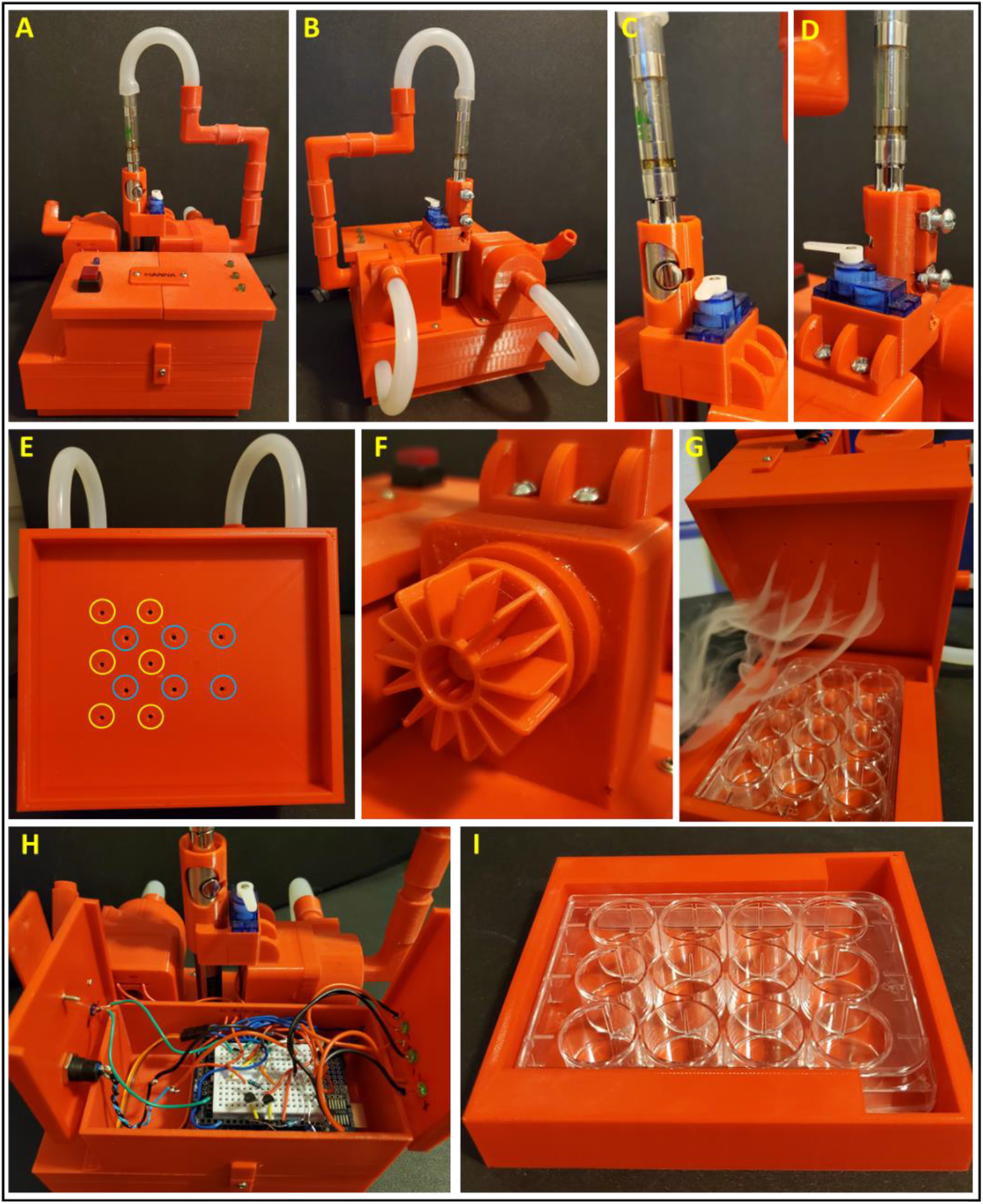
Front of vaporizing research device. B: Rear of device. C: Front of vape pen holder/button pushing mechanism. D: Reverse side of vape pen holder/button pushing mechanism. E: Inside of the sample exposure chamber, yellow circles indicate sample chamber input port, blue circles are exhale fan intake ports. F. One of two 30mm radial fans powered by a 5V DC motor. G: The device running with sample chamber open to visualize the six vapor delivery ports. H: Circuitry and Arduino Uno microcontroller. I: Bottom of sample exposure chamber which holds a 12-well culture plate.

Three-dimensional objects were drafted in Autodesk Inventor Professional 2020 (Autodesk, Inc., San Rafael, CA) and printed with PLA filament on a QIDI X-PRO 3D printer (QIDI TECH, Ruian, Zhejiang, China). Servos and fan motors are controlled by an Arduino Uno microcontroller (Boston, MA). All concepts, drafting, printing, coding, and construction was performed by laboratory personnel.

### Human Nasal Epithelial Cell (HNEC) Isolation and Expansion

HNECs were isolated through brushing of inferior and middle nasal turbinates with interdental brushes. Brushes were removed from the handle and placed into PneumaCult-Ex plus medium (StemCell Technologies inc, Vancouver, Canada). Cells were removed from the brush using a cut pipet tip. The cells were pelleted by centrifugation (200 x g for 5 minutes.) and resuspended in animal component free enzymatic digestion buffer (StemCell), briefly vortexed and then incubated at 37°C for 15 minutes. After digestion cells were centrifuged at 200 x g for 5 minutes. Cell pellets were suspended in basal cell expansion media (PneumaCult-Ex plus) and incubated at 37°C, 5% CO2. HNECs were collected and counted between 8-10 days of culture, culture media was refreshed every two days. HNECs were then either seeded directly onto transwell inserts for ALI differentiation or banked in liquid nitrogen for future experiments. Protocols for isolation of HNEC specimens from human subjects are approved by our Institutional Review Board (Study ID: Pro2019000524) and all donors provided informed consent.

### Air-Liquid Interface Differentiation

HNECs were seeded onto semi-permeable polyester tissue culture-treated inserts that were 12 mm in diameter and 0.4uM pore size and cultured in expansion media until 100% confluent (confirmed by visualization of HNECs via phase contrast microscopy). Once a confluent monolayer formed, initiation of the air-liquid interface was performed by removing media from the transwell insert and replacing the expansion media in the lower-chamber with ALI differentiation media (StemCell Technologies). At 48 hour intervals the transwell insert was rinsed with 500 μL PBS to prevent excessive mucous accumulation and the media was refreshed. Differentiation of ALI tissue was confirmed by visualization of beating cilia via phase contrast microscopy. Details of HNEC isolation, culture, expansion and differentiation are presented in Manna and Caradonna, 2021 [15].

### Vaping Sessions

Experimental vape-liquids were loaded into matching cotton wick vape cartridges operated by identical 3.4V batteries that were charged each day before use. Propylene glycol and vegetable glycerin were acquired from Florida Laboratories (Fort Lauderdale, FL), and vitamin E acetate was acquired from Lotioncrafter (Eastsound, WA). Vaping sessions began between day 20 and 25 ALI, once all tissues had differentiated. Differentiation was confirmed by visualization of beating cilia via phase contrast microscopy. Each day, the tissues were placed into the vaporizer and put through 10 inhale/exhale cycles. A single cycle depresses the button of the pen for 8 seconds. At the same time, the inhale fan turns on and runs for 15 seconds before turning off. After an additional 5 seconds the exhale fan turns on for 10 seconds to clear the experimental vapor from the chamber, ending the cycle. The inhale/exhale cycles were repeated a total of 10 times with 60 second intervals between each cycle. Our goal was not to emulate human use of a vape-pen, but to generate vapors from the pen and deliver maximal contact with experimental ALI tissues. The control tissue was placed into the experimental chamber and a vape-pen without a battery was attached for the 10 cycles. This ensured the flow of air was similar in the chamber but there was no vapor produced. All vaping sessions were performed in a biological safety cabinet.

### Human Inflammation Array

Throughout vaping experiments, the ALI media was replenished, and surfaces of tissues were washed every 48 hours, which is standard protocol for ALI tissue maintenance. For inflammatory cytokine analysis, the basal ALI media was allowed to condition for 72 hours before being collected and used to wash the apical surface of tissues. The conditioned media were centrifuged for 5 minutes at 200 x g to pellet any cells/debris. Conditioned media were then analyzed using Human Inflammation Arrays (RayBiotech AAH-INF-3-4) following the manufacturer’s protocols. Arrays were visualized using an IBright fl1500 imaging system, average pixel intensity was calculated using IBright image analysis software (ThermoFisher). Average pixel intensity from experimental groups were divided by the average pixel intensity from control groups to represent data as fold changes compared to control.

### ALI Tissue Fixation and Sectioning

ALI tissue was fixed in 10% buffered formalin at 4°C for 12-16 hours, then dehydrated via 20-minute graded alcohol (70%-80%-95%-100%) and xylene washes. Fixed and dehydrated ALI tissue was then embedded in paraffin wax blocks and 10μM sections were prepared using a microtome. For hematoxylin and eosin (H&E) staining, histological sections were deparaffinized in two changes of xylene for 5 minutes each and rehydrated via 5 min graded alcohol washes (100%- 95%-80%-70%). Sections were placed in hematoxylin for 1 minute, rinsed in tap water, dehydrated, and stained with eosin for 30 seconds. Lastly, samples were cleared with xylene before applying coverslips. Phase contrast microscopy as well as immunostaining was visualized and imaged using an ECHO Revolve fluorescent microscope.

### Immunohistochemistry

Histological sections were deparaffinized in xylene and rehydrated via 5 minute graded alcohol washes (100%- 95%-80%-70%-H_2_O). Antigen retrieval was performed in a digital pressure cooker (InstaPot) with slides submerged in antigen retrieval buffer. Sections were blocked with 5% BSA-TBST for one hour at room temperature. Primary antibodies were suspended at desired dilutions in 1% BSA-TBST and applied for one hour at room temperature. Antibodies used in this study can be found in key resources table. Secondary antibodies conjugated to either Alexa Fluor-488 or −555 were applied at 1:1000 dilution in 1% BSA-TBST for one hour at room temperature. DAPI was applied and sections were cover slipped and sealed with clear nail polish. Immunofluorescence was visualized and imaged via an ECHO Revolve fluorescent microscope running the ECHO Pro imaging application.

### Quantification of Cell Populations

Horizontal sections of continuous, undamaged ALI tissue were chosen for cell counts. Goblet cells were visualized using the periodic acid Schiff glycoprotein stain, basal progenitor populations were identified using the marker P63, and multiciliated cells were identified using the marker FoxJ1. Quantifications were performed by capturing brightfield or fluorescent images of stained 10uM-thick tissue sections and manually counting cells. Using cell counts and the length, and width of each section we determined the average cell numbers per um^2^ of tissue and extrapolated to average cell number per mm^2^ of tissue. Populations were compared via unpaired T-test; P-values <0.0001 when compared to DMSO control tissue were considered statistically significant.

## RESULTS

### VEA vapor recondenses as insoluble droplets on the mucosal surface

Unexpectedly, we noted a matte-like coating on the transwell inserts after VEA vaporizing sessions. Upon closer inspection with a phase contrast microscope, we observed the accumulation of small insoluble droplets on the mucosal surface of ALI tissues exposed to VEA vapors (Figure 2A). The relative amount of VEA droplets on the mucosal surface correlated with the amount of inhale/exhale cycles (Figure 2B). We noted that the VEA droplets were moving along with the laminar flow of the mucosal layer, albeit very slowly.

**Figure 2A.**
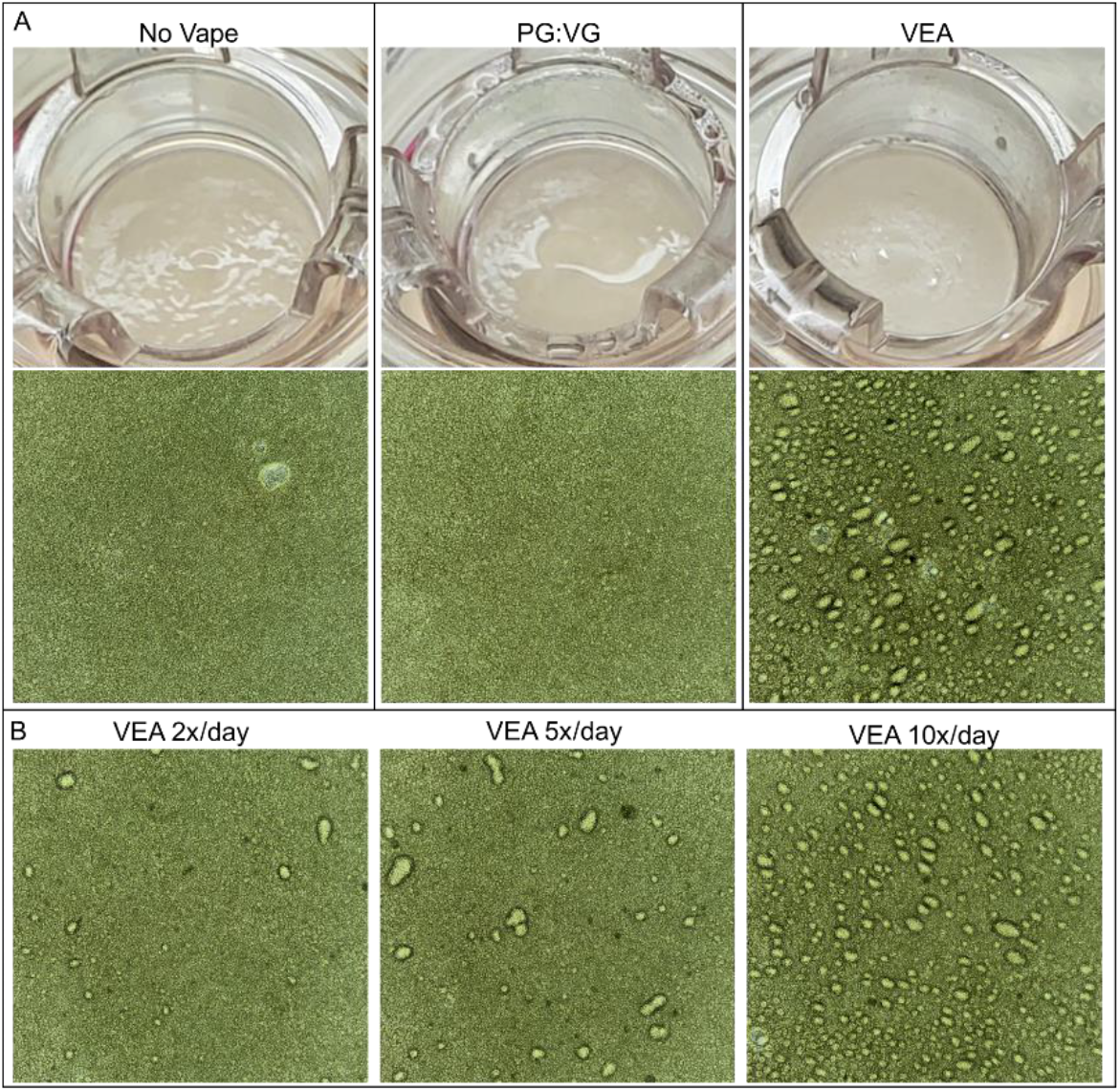
TOP: images of transwell inserts 5 minutes after 10 puffs from e-cigarettes. Inserts exposed to VEA vapor had a matte-like coating on the surface of tissues. BOTTOM: 10x phase contrast images of tissue surfaces, insoluble VEA droplets were observed on the mucosal surface after exposure to VEA vapor. B. 10x phase contrast images of tissue surfaces after 2, 5 or 10 puffs from VEA e-cigarettes. The accumulation of VEA droplets increased as exposure to VEA vapor increased.

### Tissue cultures chronically exposed to VEA vapor have increased goblet cell populations

Using the period acid Schiff (PAS) glycoprotein stain as a marker of goblet cells we found that after 2, 4, or 5-weeks of daily VEA vapor exposure the number of PAS+ goblet cells had significantly increased compared to both control and PG:VG-exposed tissue cultures (Figure 3).

**Figure 3A.**
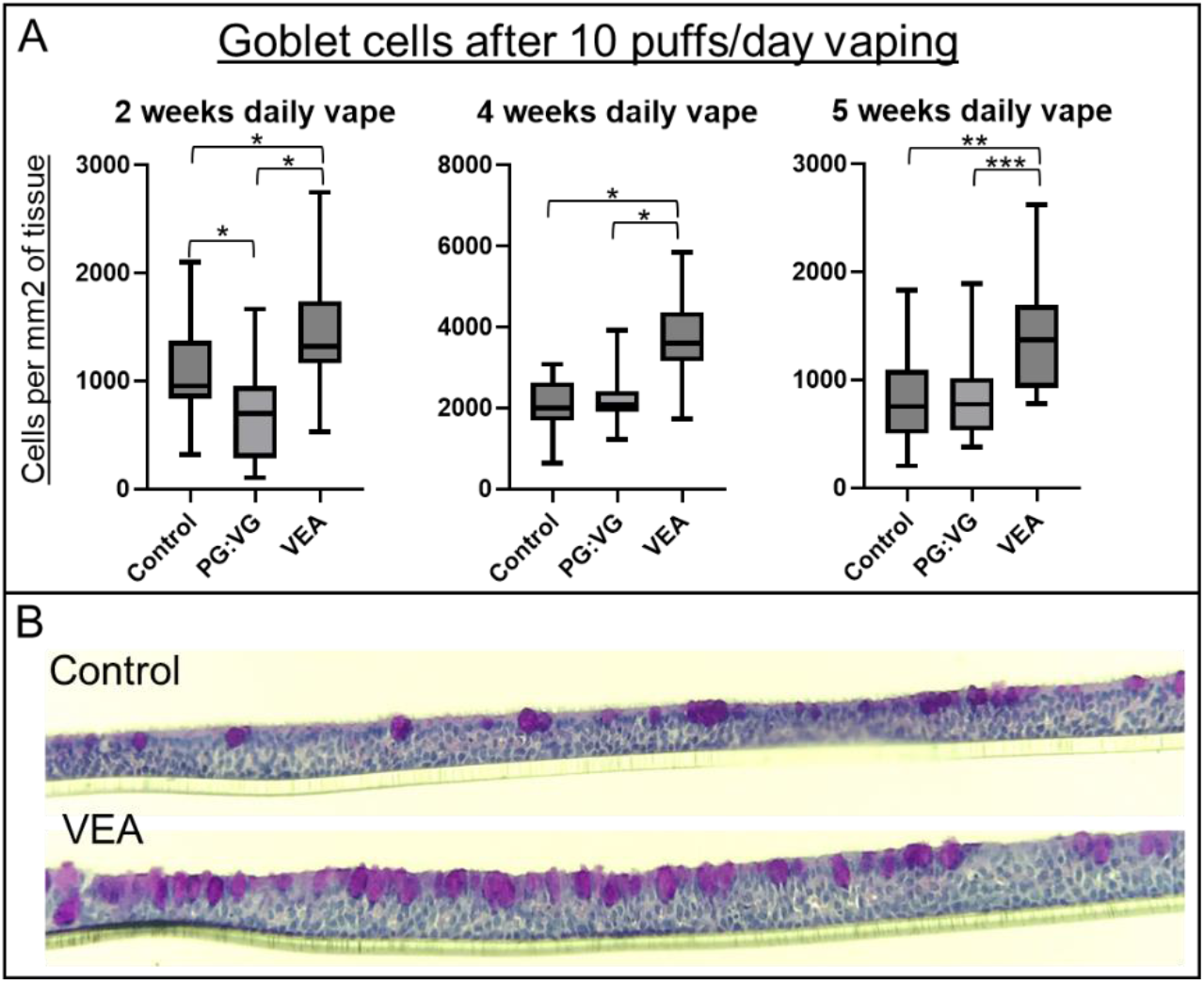
Manual goblet cell quantifications from tissue sections using Periodic Acid Schiff stain to identify goblet cells in tissues after being exposed to experimental e-cigarette vapors 10 puffs a day for 2, 4 or 5-weeks. Unpaired T-test p-values: *0.0001, **0.0002, ***0.0007. 2-week N=Ctrl:54, PGVG:29, VEA:50, 4-week N=Ctrl:39, PGVG:40, VEA:38, 5-week N=Ctrl:26, PGVG:18, VEA:14. 3B. Representative images of PAS-stained tissues after 4-weeks of daily exposure to VEA vapor demonstrating increase amount of goblet cell formation.

### Tissue cultures chronically exposed to PG:VG vapor exhibit decreases in multiciliated and progenitor cell populations

Manual quantification of multiciliated cells and progenitor cells using immunohistochemical-based markers revealed that tissue cultures chronically exposed to PG:VG vapors for 4- or 5-weeks exhibited a significant decrease in both populations when compared to control or VEA-exposed tissues (Figure 4). We did not observe this decrease in tissues exposed to PG:VG vapors for 2-weeks.

**Figure 4A:**
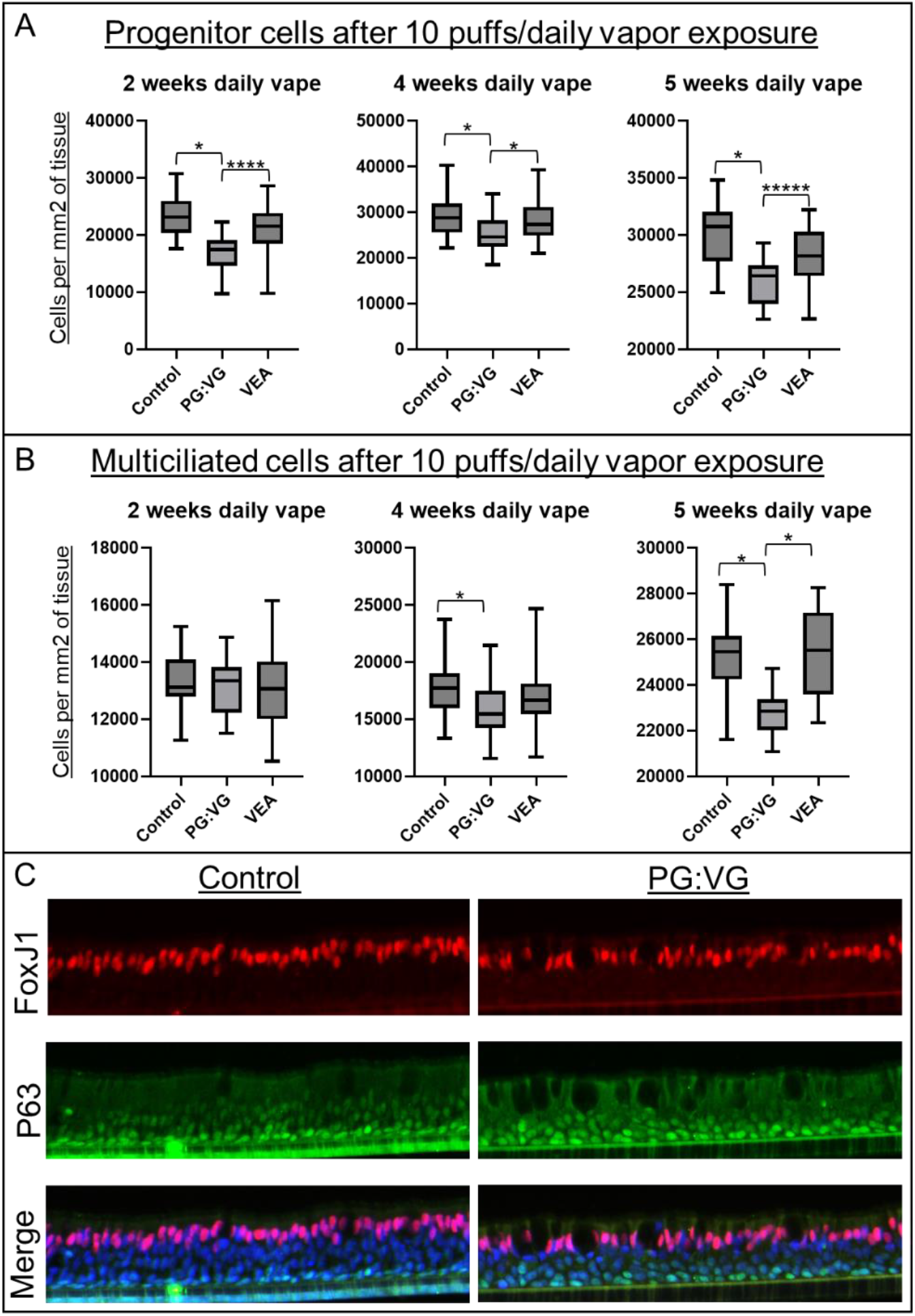
Manual progenitor cell quantifications from tissue sections using P63 stain to identify progenitor cells in tissues after being exposed to experimental e-cigarette vapors 10 puffs a day for 2, 4 or 5-weeks. 2-week N=Ctrl:32, PGVG:20, VEA:36, 4-week N=Ctrl:30, PGVG:63, VEA:59, 5-week N=Ctrl:23, PGVG:16, VEA:21. 4B: Manual multiciliated cell quantifications from tissue sections using FoxJ1 stain to identify progenitor cells in tissues after being exposed to experimental e-cigarette vapors 10 puffs a day for 2, 4 or 5-weeks. Unpaired T-test p-values: *0.0001, ****0.0046, *****0.0092. 2-week N=Ctrl:39, PGVG:21, VEA:39, 4-week N=Ctrl:60, PGVG:61, VEA:39, 5-week N=Ctrl:24, PGVG:17, VEA:23. 4C: Representative images of P63 (progenitor cells) and FoxJ1 (multiciliated cells) stained tissues after 4-weeks of daily exposure to VEA vapor displaying decreases in both progenitor cell and multiciliated cell populations.

### Chronic exposure to either PG:VG or VEA vapors caused tissue cultures to increase the secretion of immunomodulatory cytokines

At the conclusion of both the 4-week and 5-week experiments all ALI tissues exposed to PG:VG or VEA vapor demonstrated increased secretion of the same four inflammatory targets; Chemokine ligand 15 (CCL15), Interleukin-6 (IL-6), Interleukin-6 soluble receptor (IL-6sR), and Chemokine ligand 2 (CCL2). The induction of cytokine secretion was generally greater in tissues exposed to VEA vapor compared to PG:VG (Figure 5).

**Figure 5.**
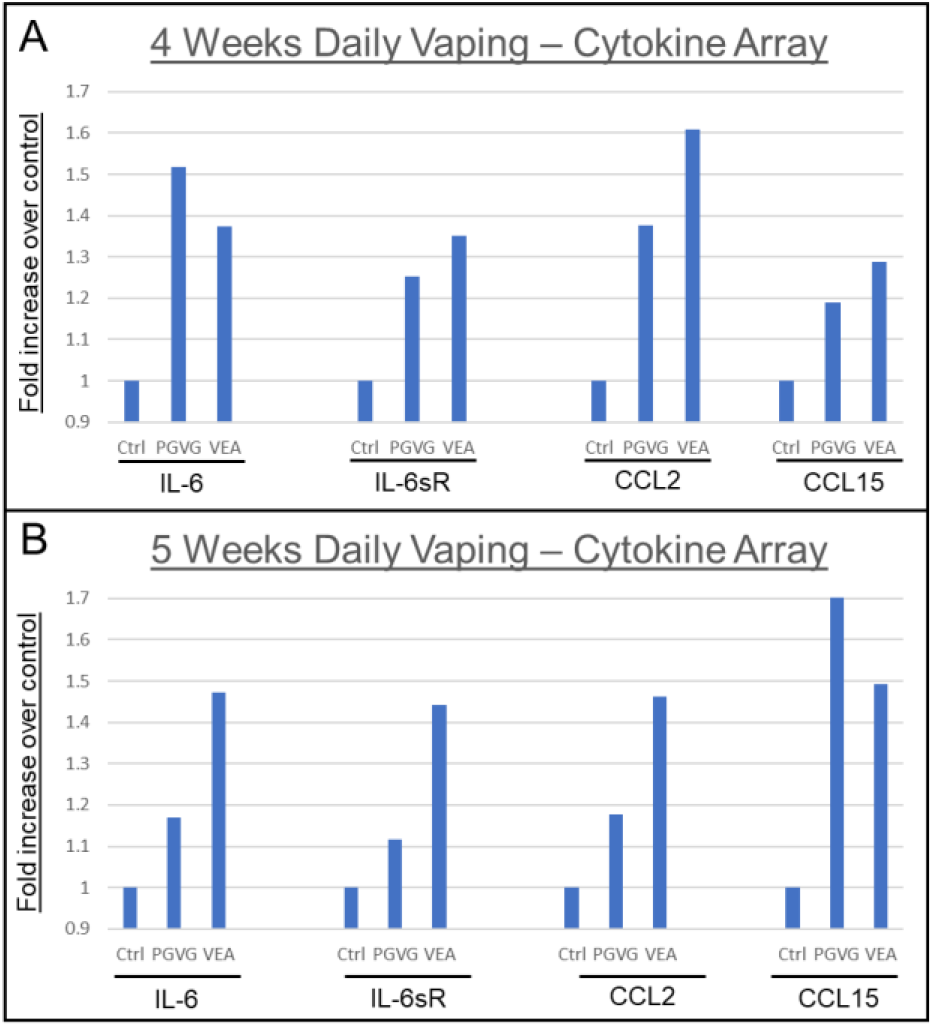
Chronic exposure to either PGVG and VEA vapors for 4 or 5 weeks resulted in an increased secretion of immunomodulatory cytokines. Human inflammation cytokine array was performed using 72-hour conditioned media combined with mucosal washes. Data represented as ‘Fold increase over control’ with control signal set to 1. Each target measured in duplicate on the array, pixel intensity calculated via iBright analysis software, the two values were used to calculate an average, PGVG and VEA average pixel intensity was divided by control avg pixel intensity for fold increase over control tissues.

## DISCUSSION

Regarding VEA, e-cigarette vapors generated from VEA recondense into oil droplets when contacting the airway mucosal surface and these droplets appear largely insoluble with airway mucous. Tissue cultures respond to the VEA debris by expanding the goblet cell compartment, most likely to produce increased amounts of mucous to clear the excess debris. A previous study in 2020 observed lipid-laden alveolar macrophages in the lungs of mice after exposure to VEA vapors for 3 hours [20]. We postulate that VEA droplets in the lung may be resistant to mucociliary clearance due to insolubility in airway mucous, and users of e-cigarettes containing VEA may accumulate VEA droplets in the lung at a faster rate than they can be cleared by immune cells. Our antibody arrays suggested that ALI tissues chronically exposed to VEA vapor increased the secretion of certain immunogenic cytokines compared to control tissues. The accumulation of VEA droplets in the lung would likely exacerbate similar inflammatory pathways and contribute to pneumonia-like symptoms observed in EVALI patients. We believe our experiments with VEA support the clinical evidence uncovered by the CDC which indicated VEA as a potential trigger of EVALI [6]. In summation, we postulate that e-cigarette vapors generated from vape-liquid containing VEA may contribute to the pathogenesis of EVALI through the accumulation of insoluble VEA droplets in the lung that behave as physical debris and provoke inflammatory responses from tissues.

We did not observe droplets or precipitates on the mucosal surface of tissues after exposure to PG:VG vapor; we hypothesize that PG:VG-vapors contain mucous-soluble compounds which enter the mucosal layer upon contact with the mucosal surface. The process of vaping PG:VG creates a multitude of toxic byproducts including aldehydes and monoxides [7] [8] [9] [10]. Some of these chemical byproducts, such as aldehydes, act as cross-linking agents which is possibly how they disrupt cellular homeostasis upon entering the mucosal layer. The tissues exposed to PG:VG vapors appeared structurally typical, yet we observed histological evidence of reduced progenitor cell and multiciliated cell populations. It is plausible that PG:VG vapor-associated toxicity increased the turnover rate of multiciliated cells via low-level crosslinking of apical cells and the progenitor cell population is being reduced from an increased demand to replenish the apical cell populations. The reduction of multiciliated cells after chronic exposure to PG:VG vapors suggests that e-cigarette users may be susceptible to the same types of injury/repair cycles that lead to airway diseases such as COPD and emphysema [18] [19] [16]. Ultimately the delineation of the acute and chronic effects of PG:VG vapor on airway tissue structure and function is a public health concern as PG:VG is the most common solvent used in nicotine-based e-cigarettes.

Currently there is a paucity of research equipment designed to operate e-cigarettes [13]. A few companies offer experimental smoking machines designed for research into the effects of smoking cigarettes. These machines are large pieces of laboratory equipment that are expensive and difficult to adapt to vape-pens and related electronic cigarettes. Our laboratory has developed a simple device that can be manufactured with minimal costs and can remain in a biological safety cabinet while still leaving surface area available for cell culture, avoiding the need for a dedicated hood. Combining this device with human airway mucociliary tissue cultures enhances the possibilities for modeling the chronic consequences of e-cigarette usage and identifying vape-liquid constituents which alter vapor toxicity. The findings from our *in vitro* study of e-cigarette vapors demonstrate the feasibility of using our research device and ALI tissues for these types of chronic airway inflammation and inhalation toxicity investigations. Using a human-derived ALI-based in vitro assay system for investigation into respiratory toxicity is an ethically sound alternative to laboratory animals that may also minimize potential non-translatable findings due to variations in the molecular pathways between humans and laboratory animals. Delivering e-cigarette vapor at a dosage below the acute destruction of tissue allows for the homeostatic maintenance of tissues in a heightened state of stress, which may allow for the presentation of tissue phenotypes which resemble alterations observed in human chronic inflammatory airway diseases [18] [19] [16]. Our research device may prove to be a simple and practical tool to leverage the assets of the ALI model system to ethically expand the research design and feasibility of human respiratory disease and inhalation toxicology investigations while reducing both the cost and space for associated studies.

